# Water-mass origin and transformation shapes seawater microbiomes in the northeastern Barents Sea and the St. Anna Trough

**DOI:** 10.64898/2026.10.16.759533

**Authors:** Olga Konovalova, Iulia Papa-Dmitrieva, Alexander Osadchiev, Anastasiia Rusanova, Elizaveta Chevokina, Daria Sibiryakina, Darya Slonova, Andrey Sobolev, Zinaida Zabudkina, Alina Demkina, Artem Isaev, Dmitry Sutormin

## Abstract

The St. Anna Trough region is a key Eurasian Arctic gateway connecting the northeastern Barents Sea, northern Kara Sea, and central Arctic Ocean. The region is characterized by complex hydrodynamics, with multiple water masses of local and Atlantic origin interacting and undergoing transformation. In this study, we investigated whether seawater microbiome composition correlates with water-mass origin and can be used to help delineate water masses. We collected 115 seawater samples representing major regional water masses - Arctic surface water, meltwater, cold dense water, Barents Sea branch water, the recently discovered Keldysh Current water, and mixed waters - and analyzed them using 16S rRNA gene amplicon sequencing. We found that microbiome composition closely followed water mass structure, which, together with thermohaline properties and depth, explained the majority (61.2%) of microbiome composition variation. Distinct microbial patterns were associated with Atlantic and Arctic water masses, except for the Arctic surface water, which lacked a distinct microbiome and resembled those of meltwater or the Keldysh Current microbiomes. Differential abundance and network analyses revealed characteristic bacterial markers: SAR11 (Clades Ia and IV), *Roseibacillus*, SV1-3 (*Microbacteriaceae*), SAR86 clade, and AEGEAN-169 marine group in the Keldysh Current; *Halieaceae*, *Planktomarina*, NS5 marine group, and *Sulfitobacter* in meltwater; SUP05 cluster, SAR86, SAR92, and SAR324 clade in cold dense water; *Nitrincolaceae* and *Amylibacter* in Barents Sea branch water; *Flavobacteriaceae* and Chitinophagales in mixed waters. These differences were driven primarily by shifts in relative abundance of widespread Arctic bacterial taxa rather than by differential prevalence of bacterial groups. Our results demonstrate that seawater microbiomes can serve as sensitive biological tracers of water mass origin and transformation in the Eurasian Arctic. Furthermore, the distinct Keldysh Current microbiome supports its recognition as a novel hydrographic feature and indicates its role as a biological corridor transporting Atlantic-associated biota eastward toward the Kara Sea. These findings suggest that “Atlantification” of Eurasian Arctic includes a microbial component, highlighting marine microbiomes as indicators of ongoing ecological changes in the Arctic Ocean.

## Introduction

The Arctic Ocean is considered a global carbon sink, contributing to the worldwide carbon cycle (MacGilchrist et al., 2014). The region is strongly impacted by global warming, with a temperature increase rate estimated to be four times faster than in the rest of the world (Rantanen et al., 2022). Microcosm experiments and field studies predicted that warming and increasing CO_2_ can lead to a decrease in phytoplankton biomass and enhancement of bacterial production in Arctic water (Vaque et al., 2019; Boras et al., 2010). Such changes may significantly impact the global carbon cycle and create a positive feedback loop that could exacerbate global warming in the future.

The inflow of Atlantic water masses significantly contributes to Arctic warming. The inflow, enhanced during the historic era, leads to sea-ice decline and weaker water stratification, resulting in rapid Arctic “Atlantification” (Polyakov et al., 2017; Tesi et al., 2021). The warm, saline, and nutrient-rich inflow from the Atlantic strongly affects the ecosystems of the Arctic Ocean, a phenomenon called “biological Atlantification” (Csapó, Grabowski, Węsławski, 2021; Polyakov et al., 2020; Polyakov et al., 2025).

Atlantic water mass enters the Arctic Ocean through the Fram Strait (the northern or the Fram Strait branch hereafter referred to as FSBW), a passage between Greenland and Svalbard, and the Barents Sea (the southern or the Barents Sea branch hereafter referred to as BSBW) (Rudels et al., 2013; Schauer et al., 2004) (**Figure 1**). The effects of Atlantic influx on water microbiomes were extensively studied in the Fram Strait region and the Norwegian Sea, and the obtained results overall support the concept of “biological Atlantification” from a microbiome perspective (Priest et al., 2023; Carter-Gates et al., 2020; Sinha et. al., 2017; Fadeev et al., 2018; Wietz et al., 2023). Concordantly, a pioneering study correlating microbiome composition with water masses genesis in Eurasian and Canada Basins showed that Atlantic water has a distinct microbiome (Galand et al., 2010). Similar results were obtained in the Central Arctic Ocean for the Chukchi Borderland (Han et al., 2015) and at the Beaufort Gyre (Vipindas et al., 2023), indicating that Atlantic water masses transport Atlantic microbiota over long distances of thousands of kilometers and/or Atlantic water masses induce microbiome changes on their path. Further, these observations suggested the identification of Arctic water masses based on microbiome composition (Han et al., 2015).

**Figure 1.**
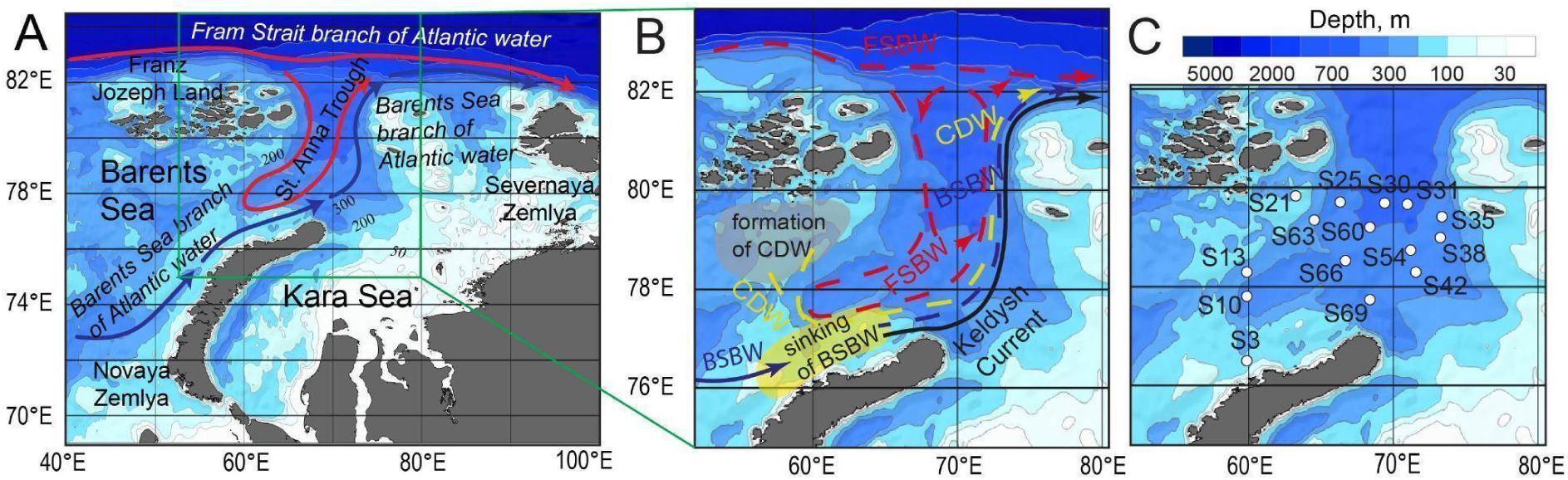
Geography and macro-/meso-scale hydrophysical features of the study area. (**A**) Flow of the Fram Strait (FSBW, red arrows) and Barents Sea (BSBW, blue arrows) branches of Atlantic-origin water in the northeastern part of the Barents Sea and the northern part of the Kara Sea. (**B**) Scheme of currents in the northeastern part of the Barents Sea and the northern part of the Kara Sea. Solid lines indicate surface currents; dashed lines indicate intermediate and bottom currents. CDW indicates cold dense water. (**C**) Geographical location of hydrographic and water sample stations in the study area during the oceanographic survey performed on September 11-30^th^ 2023.

Hydrographic data indicates that BSBW propagates to the East through the Barents Sea to the Kara Sea and then turns to the north at the St. Anna Trough, where it meets FSBW that makes a side counterclockwise loop in this region (Schauer et al., 2004; Osadchiev et al., 2022) (**Figure 1**). The convergence of BSBW and FSBW Atlantic branches may suggest a strong impact of Atlantic inflow and progressive “Atlantification”. However, microbiome studies of the St. Anna Trough region remain scarce, which limits our understanding of ecological processes in this area (Csapó, Grabowski, Węsławski, 2021). Bacterioplankton abundance and its biomass have been assessed in the Kara Sea (Sazhin et al., 2010), but community composition has been reported only for a few samples obtained from the surface layer (Bezzubova et al., 2024). A recent analysis of surface water microbiomes in the northeastern Barents Sea, west of the St. Anna Trough, revealed two distinct microbiome clusters in the surface water, but they were not associated with specific water masses or Atlantic inflow (Namsaraev et al., 2023).

To fill this gap, we performed a systematic collection of surface and subsurface seawater (up to 50 m depth) in the St. Anna Trough area and adjacent northeast region of the Barents Sea. We delineated water masses in this region based on their hydrographic and hydrophysical properties and complemented this analysis with 16S rRNA gene metabarcoding to reveal the compositions of seawater microbiomes. We found that different water masses have distinct microbiome compositions, and that their genesis may be predicted based on microbiome information. Our study indicates the ecological influence of Atlantic inflow in the St. Anna Trough region and paves the way for monitoring of the “biological Atlantification” process in this region.

## Materials and Methods

### Hydrology of the study area

The main circulation feature and water masses in the northeastern part of the Barents Sea and the St. Anna Trough are shown in **Figure 1A**. BSBW goes through the Barents Sea and experiences substantial cooling along its route due to year-round absence of sea ice in the southwestern part of the Barents Sea (Rudels et al., 2004; Smedsrud et al., 2013). At the northeastern part of the Barents Sea, BSBW becomes dense enough to lose contact with atmosphere (Parsons et al., 1996; Schauer et al., 2002). It transforms into a bottom-advected current and flows further to the north through the St. Anna Trough located in the northern part of the Kara Sea (Lien and Trofimov, 2013; Dmitrenko et al., 2015). A share of FSBW also inflows to the St. Anna Trough, albeit from the continental slope located to the north (Schauer et al., 2002; Lind and Ingvaldsen, 2012). FSBW recirculates in the St. Anna Trough where it interacts with other local waters (including BSBW) and becomes colder and fresher (Osadchiev et al., 2022; Zabudkina et al., 2025).

Surface layer in the northeastern part of the Barents Sea and the St. Anna Trough is occupied by Arctic surface water (ASW), which is the result of mixing sea ice meltwater with saline seawater (Oziel et al., 2016). Arctic surface water interacts with the northern periphery of BSBW flow before it loses contact with the atmosphere. As a result, the northern periphery of BSBW becomes fresher and remains at sea surface after the major volume of BSBW has sunk. This mixture of BSBW and ASW forms the Keldysh Current, the warm and surface-advected current that follows the bottom-advected flow of BSBW. The Keldysh Current serves as a fast transit zone along the northern coast of Novaya Zemlya, the eastern flank of the St. Anna Trough and the continental slope eastward from the St. Anna Trough (Osadchiev et al., 2026). On the other hand, the Keldysh Current isolates surface water at the shallow southern part of the Kara Sea dominated by river discharge and surface water at the central and northern part of the Barents Sea (Osadchiev et al., 2023; Savin and Osadchiev, 2025).

In winter, surface-advected water masses experience intense cooling at multiple shallow banks located either in the ice-free part of the Barents Sea, or at areas with intense polynya activity (Arthun et al., 2011; Ivanov et al., 2021). It results in the formation of cold dense water (CDW) with different thermohaline properties, which depend on thermohaline properties of the initial surface water mass, i.e., ASW or BSBW (if it yet did not lose contact with atmosphere) (Lien and Trofimov, 2013; Ivanov et al., 2020). In summer, sea ice melting results in formation of short-lived (several weeks) low-saline meltwater lenses at sea surface (Osadchiev et al., 2024).

In summary, the northeastern part of the Barents Sea and the St. Anna Trough is composed from the following water masses: Arctic surface water (ASW), meltwater (MW), Barents Sea branch water (BSBW), Fram Strait branch water (FSBW), Keldysh Current water (KC), cold dense water (CDW). All these water masses, except ASW and MW, form distinct currents at the study area, which flow off the St. Anna Trough and the eastward along the continental slope (**Figure 1B**).

### Sample collection and processing

Seawater samples were collected in the northeastern part of the Barents Sea and the St. Anna Trough on 11-30 September 2023 onboard the research vessel “Dalnie Zelentsy” (**Figure 1C**). Water samples were collected in 4 replicates from the sea surface and sub-surface (30 and 50 m depth) layers at 15 sampling stations (**Supplementary Table 1**). At all of 15 sampling stations, surface-to-bottom thermohaline measurements were also performed using an AML-3 LGR CTD instrument. For water collection, a Niskin sampler (5 L) was manually deployed. 1 L of a water sample was immediately filtered through the 0.22 μm Sterivex filter (Merck-Millipore) in a ship laboratory. All filters were stored at −20°C and then transferred to the Skoltech laboratory (Moscow, Russia) for DNA isolation. A contamination control was prepared by filtering 1 L of sterile laboratory water through 0.22 μm Sterivex filter (Merck-Millipore). DNA extraction was performed as previously described (Demkina et al., 2023). Briefly, the filter membrane was removed from the plastic cover and cut into small pieces. Membrane fragments were incubated in 200 μL of STET lysis buffer (50 mM Tris-HCl, pH 8.0; 50 mM EDTA; 5% Triton X-100; 200 mM NaCl; supplemented with fresh 10 mg/mL lysozyme) at 37°C for 1 h with shaking at 600 rpm. Upon incubation, lysates were collected and DNA was extracted using the DNeasy Blood and Tissue Kit (Qiagen), following the manufacturer protocol. 115 DNA samples collected from 15 sampling stations (29 sampled water layers in total) were used for further analysis.

### 16S rRNA gene V3-V4 region amplicon sequencing and data analysis

Amplicon library preparation was performed according to the Illumina guide (https://support.illumina.com/documents/documentation/chemistry_documentation/16s/16s-metagenomic-library-prep-guide-15044223-b.pdf, 03.04.2026) and using the 341F (TCGTCGGCAGCGTCAGATGTGTATAAGAGACAGCCTACGGGNGGCWGCAG) and 785R (GTCTCGTGGGCTCGGAGATGTGTATAAGAGACAGGACTACHVGGGTATCTAATCC) primers for the V3-V4 region of the 16S rRNA gene. The amplicon libraries were barcoded, pooled as a single sequencing batch, and sequenced using Illumina NovaSeq 6000 with 2 × 250 bp paired-end layout at Evrogen LLC (Moscow, Russia). Analysis of 16S gene amplicon sequencing data was performed using a previously developed Snakemake pipeline (https://github.com/chorzow/16S_PE). Briefly, the initial quality of sequencing data was checked using FastQC (https://www.bioinformatics.babraham.ac.uk/projects/fastqc/, 03.04.2026) and then reads were trimmed and filtered using Trimmomatic v. 0.39 (PE -phred33 ILLUMINACLIP:2:30:10 LEADING:0 TRAILING:0 SLIDINGWINDOW:4:15 HEADCROP:17 MINLEN:150) (Bolger, Lohse, Usadel, 2014). Survived read pairs were used for further analysis. Reads that passed quality control were processed with the DADA2 pipeline v. 1.14.1 (Callahan et al., 2015). Resultant amplicon sequence variants (ASVs) were clustered using MMseqs2 v. 1.79 (coverage > 0.8, identity > 0.98) (Steinegger, Söding, 2017) and the obtained cluster representative sequences were further used as operational taxonomic units (OTUs). OTUs were imported into DADA2 for taxonomy assignment using the SILVA SSU database v.13835 (Quast et al., 2013). Scarce OTUs with an average relative abundance of <0.001% were removed. Samples with a high relative abundance of *Pseudoalteromonas* (>10%) were excluded from further analysis. Alpha-diversity (Chao1 and Shannon indices), beta-diversity (Bray-Curtis dissimilarity calculation, principal coordinate analysis, PERMANOVA), and taxonomic analyses were performed using R packages phyloseq v. 1.48.0, ggplot2 v. 3.5.1, dplyr v. 1.1.4, and vegan v. 2.6.8. Differential abundance analysis was performed using LefSe from the microbiomeMarker R package v. 1.4.0 (Segata et al., 2011; Cao et al., 2022). Microbial abundance correlation networks were built using SpeSpeNet v. 1.0 (van Eijnatten et al., 2025). Marker taxa were selected using the following filtration rules. Taxa were excluded if: (i) their OTUs were enriched in more than one water mass according to LefSe; (ii) their fold change (FC) of the max value to the next-highest water mass value was below 1.3, (iii) their relative abundance in the corresponding water mass was below 1%, or (iv) the coefficient of variation (CV) of relative abundance across water masses was below 0.5. After the filtration step, specificity of the remaining taxa was assessed by calculating the rank-biserial effect size. For each water mass, the taxon with the highest mean relative abundance was selected as the first marker; the second marker was then chosen by comparing the highest effect sizes of the remaining candidates, retaining the more abundant taxon if the effect size difference was less than 0.01, and the taxon with the higher effect size otherwise.

## Results

### Delineation of water masses based on thermohaline properties and geographical location of seawater samples

We performed a systematic collection of surface and subsurface seawater (up to 50 m depth) from 15 sampling stations in the St. Anna Trough area and adjacent northeast region of the Barents Sea in September 2023. To link seawater samples collected in this study with water masses and determine their origin, we first analyzed their thermohaline properties. We found that the temperature and salinity of the 29 analyzed water layers were sufficient to delineate the main surface and subsurface water masses in the study area (**Figure 2**).

**Figure 2.**
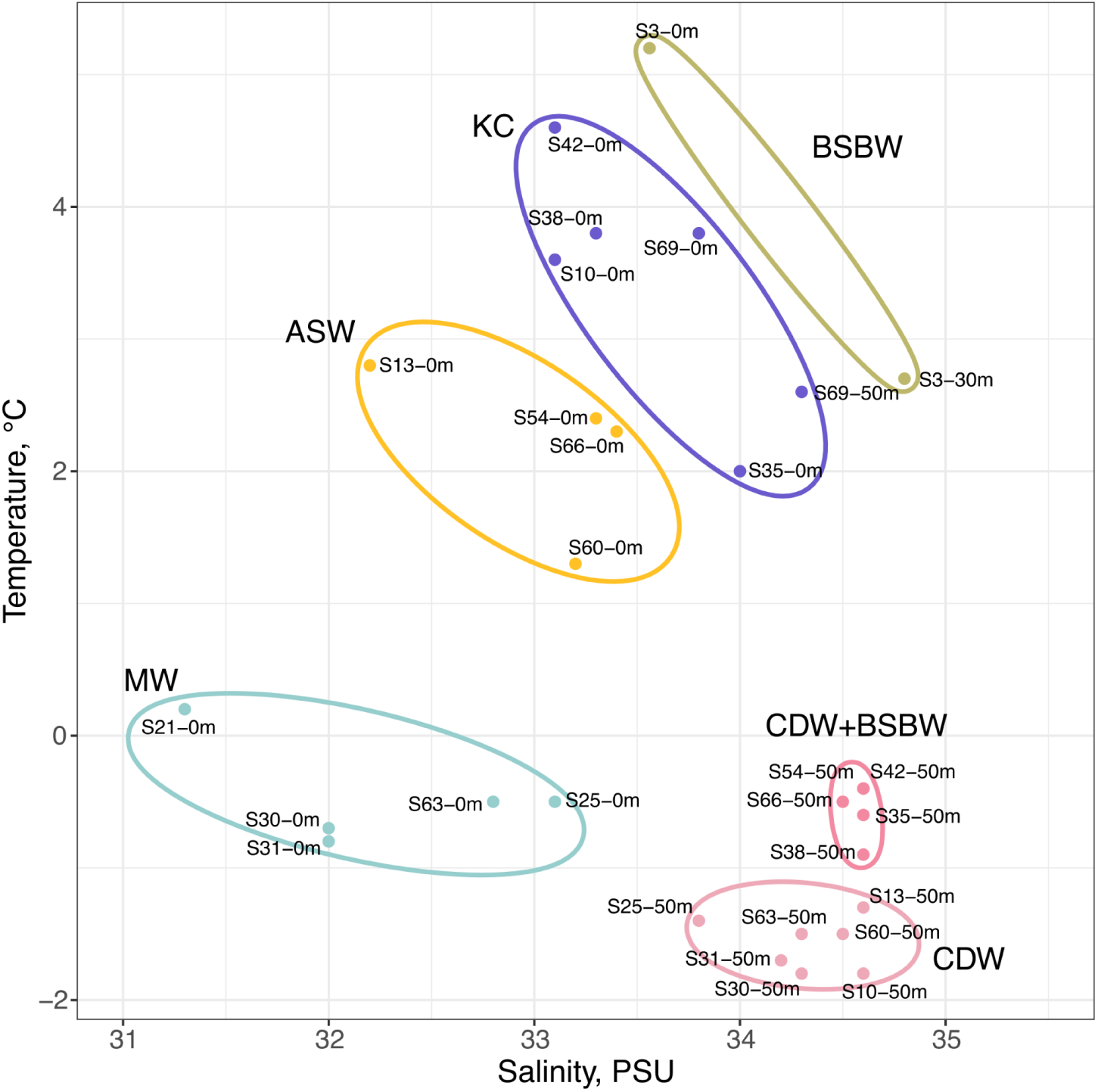
Classification of water samples collected from the northeastern Barents Sea and the St. Anna Trough region into water masses based on thermohaline properties. TS-diagram of 29 surface and subsurface water layers collected in the northeastern part of the Barents Sea and the St. Anna Trough in September 2023. Colored ellipses indicate water masses, namely, meltwater (MW) (cyan), Arctic surface water (ASW) (yellow), Keldysh Current (KC) (violet), Barents Sea branch water (BSBW) (green), cold dense water (CDW) (light pink), cold dense water mixed with Barents Sea branch water (CDW + BSBW) (dark pink).

Decreased salinities (31 to 33.1 psu) and temperatures (−1 to 0.1 °C) were registered in surface samples at stations S21, S25, S30, S31, S63, clearly indicating their meltwater (MW) origin. Station S3 demonstrated both increased temperature (5.2 °C) and salinity (33.5 psu) among the surface samples, which is typical for BSBW. Other surface samples were more closely located at the temperature-salinity (TS) diagram. However, they could be divided into two groups, namely, colder and fresher ASW (stations S13, S54, S60, and S66) and warmer and more saline KC (stations S10, S35, S38, S42, S69).

Geographical location of the analyzed stations and identified water masses were consistent with the general surface circulation scheme in the study area (**Figure 1C**). Specifically, station S3 (BSBW) was located in the area where BSBW was shown to have contact with the atmosphere. Station S10 (KC) was located at the northern periphery of BSBW, which further transforms into KC and flows along the southeastern and eastern flank of the St. Anna Trough, where stations S35, S38, S42, and S69 were sampled (all KC). Stations S13, S54, S60, and S66 (all ASW) were located northward from BSBW and KC flow, where presence of ASW is expected. Further northward, surface layer is occupied by MW, which, at the time of the oceanographic survey, had not yet dissipated following the recent sea-ice retreat that occurred two to three weeks earlier. Consistently, presence of MW is observed at the most northern stations - S21, S25, S30, S31, and S63.

At the subsurface layer, station S3 had increased salinity (34.8 psu) and temperature (2.7 °C) that links it to BSBW. All other subsurface samples had similar thermohaline properties, however, warmer (−1 to 0 °C) and more saline (∼34.5 psu) water was registered in the southeastern part of the study area at stations S35, S38, S42, S54, and S66. This geographical area corresponds to the inflow of Atlantic-origin waters, namely, bottom-advected BSBW and surface-advected KC. As a result, we associated this subsurface water mass to a mixture of CDW and BSBW. Colder (−2 to −1 °C) and less saline (33.8 to 34.5 psu) water was observed at subsurface layers at stations S10, S13, S25, S30, S31, S60, and S63. Based on thermohaline properties, geographical location, and hydrological model of this area, we linked these samples to CDW formed in winter at polynyas near Franz Joseph Land.

Taken together, using thermohaline properties of water samples and by considering the geographical location of sampling stations and hydrological model of the studied region, we associated water samples with known water masses.

### Microbiome composition clustering generally recapitulates the water mass origin

Next, we profiled seawater microbiomes from the sampling stations using the 16S rRNA gene amplicon sequencing (**Supplementary Table 2**). The generated 16S datasets were sequenced to saturation, as indicated by rarefaction curves (**Supplementary Figure 1A**-**B**), and had low contamination levels (**Supplementary Figure 1C**-**D**), thus, adequately capturing the diversity of seawater microbiomes.

We employed beta-diversity analysis to compare the microbiome composition in different water masses. Initially, using principal coordinates analysis (PCoA) based on Bray–Curtis dissimilarity, we found that some samples enriched with *Pseudoalteromonas* were clustered separately or scattered (**Supplementary Figure 2A**). We also noticed that enrichment of *Pseudoalteromonas* was inconsistent between replicates (**Supplementary Figure 3**). This might reflect occasional capture of plankton or particulate matter, as surface-associated microbiomes are known to be enriched in this taxon (Holmström, Kjelleberg, 1999; Fontanez et al., 2015; Smriga et al., 2016). We reasoned to exclude outlier samples with a relative abundance of *Pseudoalteromonas* ≥10% and performed alpha- and beta-diversity analysis on remaining 104 out of 115 microbiomes. Alpha-diversity analysis revealed that CDW and CDW+BSBW water masses had increased diversity, while the diversity of BSBW and ASW was decreased (**Supplementary Figure 2B**). With beta-diversity analysis, we found that samples obtained from different water masses, except ASW, tend to form separate clusters (PERMANOVA, p < 0.001) (**Figure 3A**).

**Figure 3.**
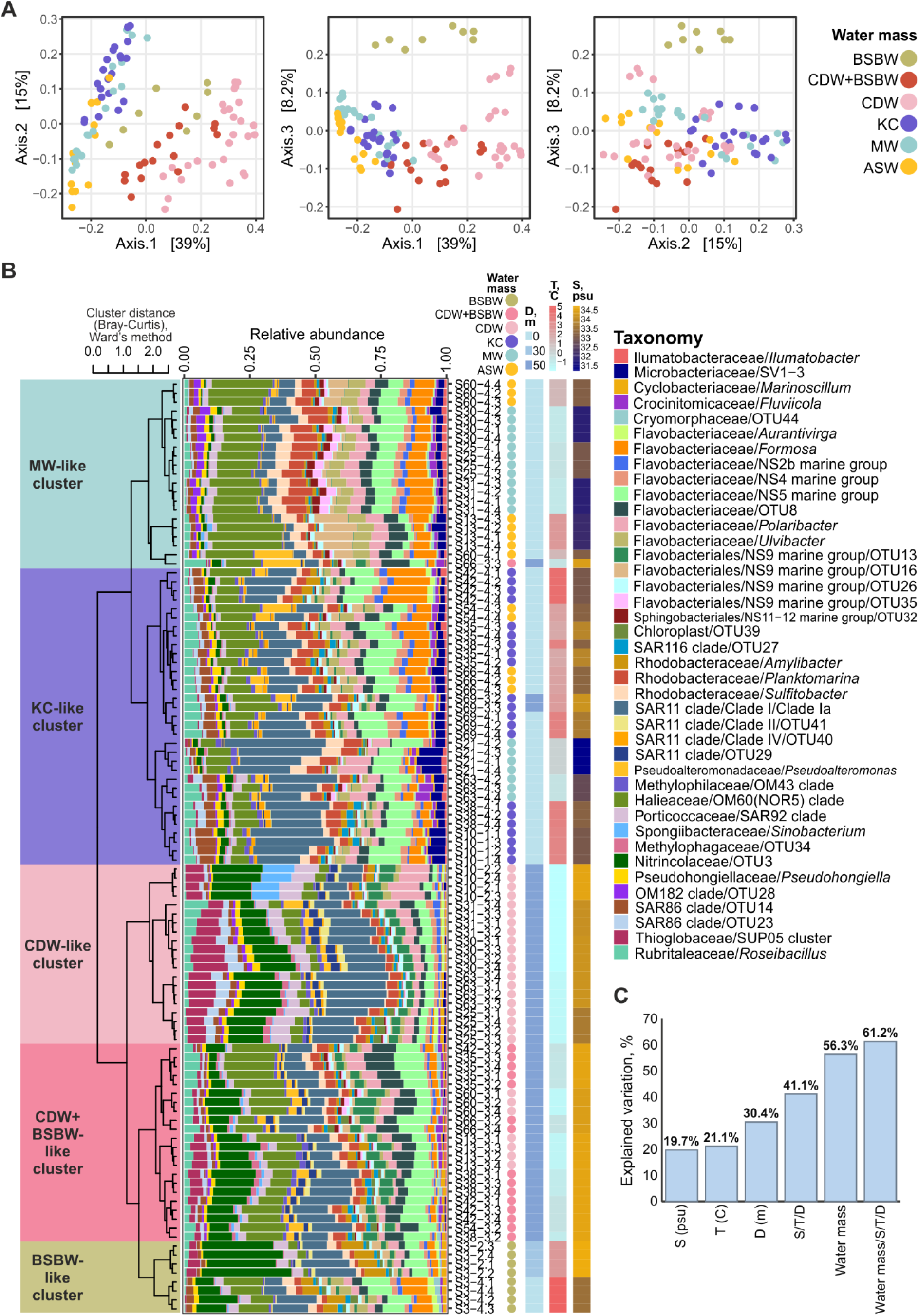
Diversity of seawater microbiomes in St. Anna Trough region is explained by environmental parameters and water mass type. (**A**) Beta-diversity (PCoA based on Bray-Curtis dissimilarity) for collected seawater samples. Samples are colored according to the water mass type. (**B**) Relative abundance of top-40 bacterial genera (or OTUs, if a genus was not determined) in the studied seawater samples. Samples are clustered by Ward’s method based on Bray-Curtis dissimilarity. The dendrogram on the left reflects hierarchical clustering of microbiomes. Heatmaps on the right represent sampling depth, temperature, and salinity at the corresponding collection sites and horizons. (**C**) Variation in microbiome composition, explained by environmental factors (temperature – T, salinity – S, depth – D), water mass types, and their combinations.

Next, we performed unsupervised clustering on the dataset. This analysis revealed five distinct groups (PERMANOVA, p < 0.001) (**Figure 3B**). MW samples dominated the first cluster, which was therefore designated MW-like; it also included several ASW samples. All KC samples, including those collected at 50 m depth from station S69 and identified as KC based on thermohaline properties, were grouped within the second cluster, thus designated as KC-like. This cluster also contained several ASW and MW samples. The third and fourth clusters consisted mainly of CDW and CDW+BSBW samples, respectively, and thus were designated as CDW-like and CDW+BSBW. The fifth cluster comprised BSBW samples from station S3 (0 and 30 m) and was designated BSBW-like.

Overall, we observed a good correspondence between MW, KC, CDW, CDW+BSBW, and BSBW water masses and microbiome clustering (**Figure 4**). Interestingly, ASW lacked a distinct microbiome and was represented by KC-like and MW-like microbiomes.

**Figure 4.**
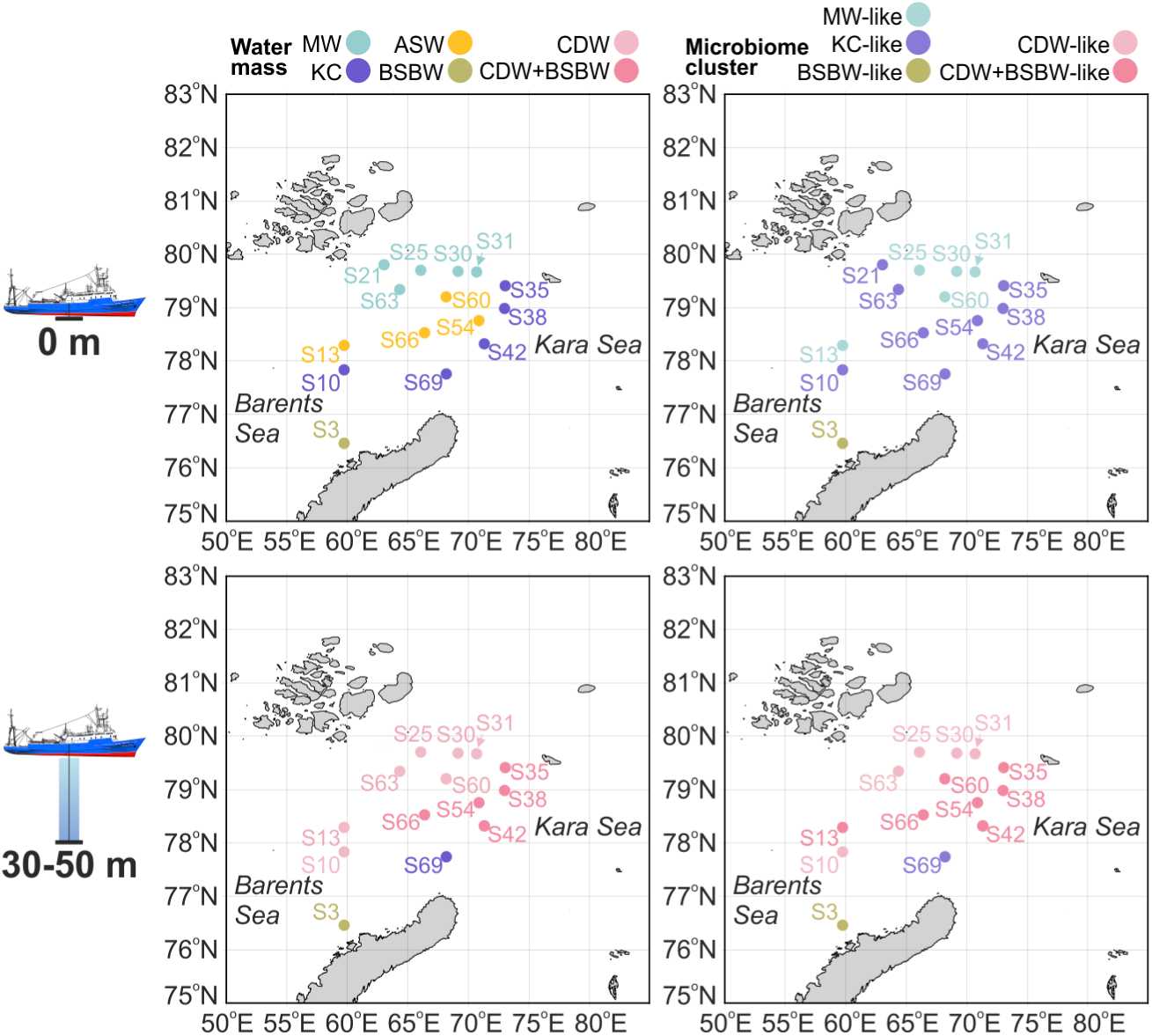
Correspondence between water masses and microbiome clusters. Maps on the left and on the right show the assignment of collection sites to water masses and microbiome clusters, respectively. Maps on the top and the bottom represent surface (0 m) and subsurface (30-50 m) samples, respectively.

### Environmental parameters and water mass origin explain most of the microbiome composition variation

To elaborate on the apparently direct correspondence between microbiome clustering and water masses origins, we next explored the effects of environmental variables on microbiome composition.

Using dbRDA stepwise significance testing, we found that salinity explained 19.7% of the variation (R^2^ adjusted=0.189, ANOVA p-value <0.001) in bacterial community dissimilarity, while sampling depth and temperature explained, respectively, 30.4% (R^2^ adjusted=0.297, ANOVA p-value <0.001) and 21.1% (R^2^ adjusted=0.203, ANOVA p-value <0.001) of the variation (**Figure 3C**). A combination of these three environmental parameters explained 41.1% of the variation (R^2^ adjusted=0.393, ANOVA p-value <0.001). The water genesis (water mass type) contributed to 56.3% (R^2^ adjusted=0.541, ANOVA p-value <0.001) of the variation.

A combination of these four environmental variables, while accounting for potential collinearity between environmental parameters (salinity, temperature, and depth) and water mass types, explained a total of 61.2% of the microbiome composition variation (R^2^ adjusted=0.579, ANOVA p-value <0.001).

We concluded that water mass origin, salinity, temperature, and depth explained the majority of microbiome composition variation.

### Marker bacterial taxa of water masses from the St. Anna Trough region

To align the microbiome clusters with the water mass categories and identify marker bacterial taxa, we selected samples from representative stations where the two classifications matched (e.g., MW samples clustering as MW-like) for further analysis and designated these as core water masses. The core water masses had an overall similar prevalence of OTUs, with a small fraction of mass-specific OTUs, indicating that OTU frequency rather than presence/absence drives the differences between them (**Figure 5A**). Differential abundance analysis using LefSe revealed that Clade Ia (SAR11), *Roseibacillus* (*Rubritaleaceae*), SV1-3 (*Microbacteriaceae*), and the SAR86 clade were the top bacterial taxa associated with KC core, while OM60 (NOR5) clade (*Halieaceae*), *Formosa* (*Flavobacteriaceae*), *Planktomarina* (*Rhodobacteraceae*), and NS5 marine group OTU4 (*Flavobacteriaceae*) were top-enriched in MW core (**Figure 5B**). Different *Flavobacteriaceae* (unassigned OTU8, NS9 marine group, and NS5 marine group OTU20) were enriched in CDW+BSBW core, and SUP05 cluster (*Thioglobaceae*), SAR86 (unassigned OTU12), and SAR92 (*Porticoccaceae*) were enriched in CDW core. BSBW core was enriched with *Nitrincolaceae* (unassigned OTU3), *Amylibacter* (*Rhodobacteraceae*), and ML602J-51 (*Microbacteriaceae*).

**Figure 5.**
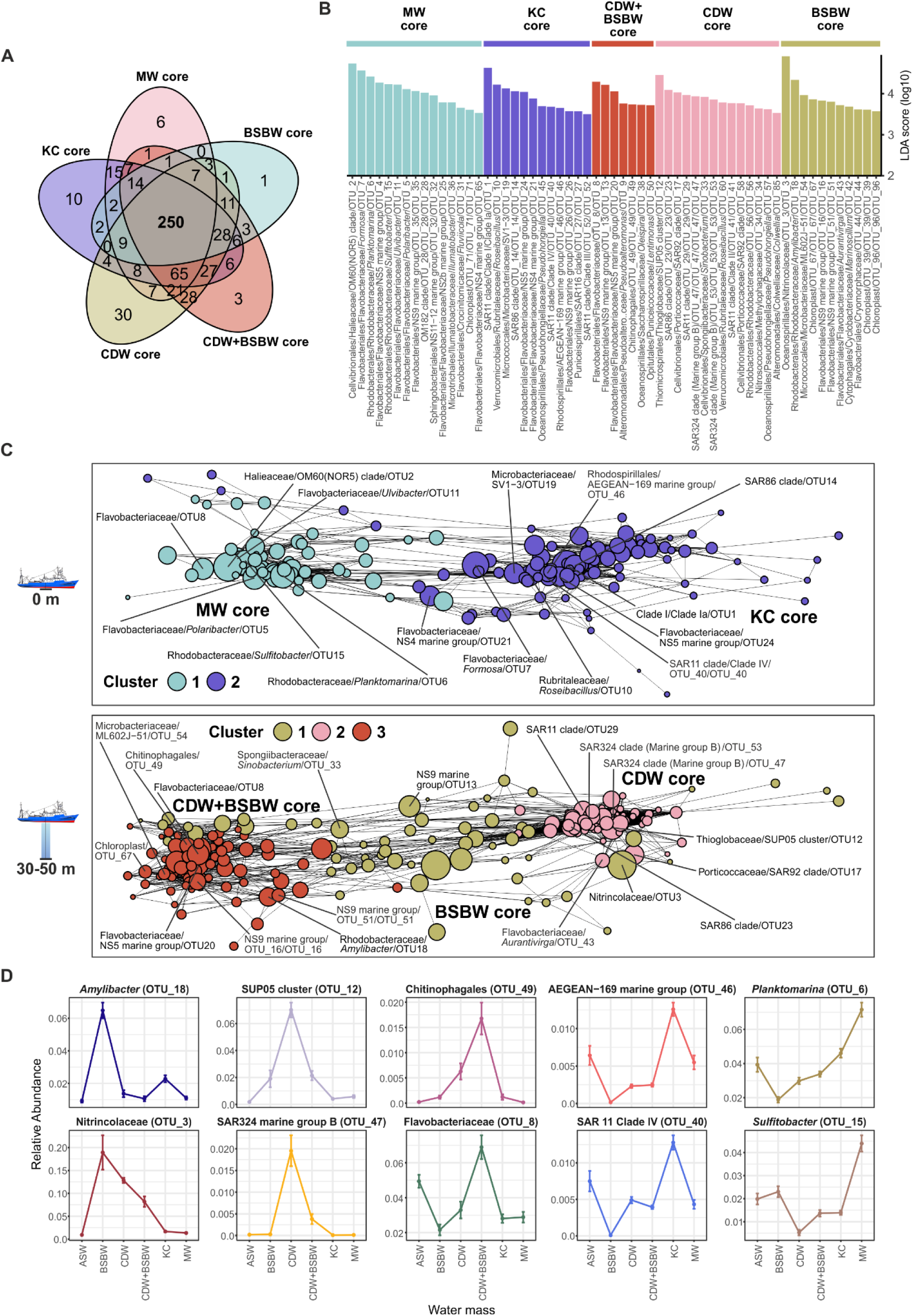
Identification of bacterial marker taxa specific for St. Anna Trough surface and subsurface water masses. (**A**) A Venn diagram showing overlaps between sets of OTUs present in core water mass samples. Numbers in the intersection indicate the number of OTUs. (**B**) LDA score barplot for top differentially abundant OTUs (LDA score>3.5) identified with LefSe for core water mass sample groups. (**C**) OTU correlation network for surface (top) and subsurface (bottom) core water mass samples obtained with SpeSpeNet with default conditions. Differentially abundant OTUs with the highest LDA scores (as calculated by LefSe) are indicated. (**D**) Relative abundance of selected marker bacterial taxa across water masses.

Next, we performed a correlation network analysis that recapitulated the existence of global surface and subsurface microbiomes and again highlighted a set of OTUs associated with *Pseudoalteromonas* (presumable plankton/particulate matter microbiome) (**Supplementary Figure 4**, **Supplementary Table 3**). The same analysis of surface samples recapitulated two robust network clusters corresponding to the KC and MW core groups: all top-enriched OTUs except *Formosa* (*Flavobacteriaceae*) clustered within their respective network clusters (**Figure 5C**, top; **Supplementary Table 4**). A network for subsurface samples revealed two robust clusters corresponding to CDW and CDW+BSBW core groups, and a loose cluster for the BSBW (**Figure 5C**, bottom; **Supplementary Table 5**). The CDW+BSBW network cluster contained all corresponding top-enriched OTUs except NS9 marine group (*Flavobacteriaceae*), and the CDW cluster contained all top-enriched OTUs. In contrast, the BSBW-enriched taxa were mostly clustered within CDW and CDW+BSBW network clusters with only the *Nitrincolaceae* (unassigned OTU3) designating the BSBW network cluster.

Finally, by selecting the taxa showing an agreement between the enrichment and correlation analyses and further constraining this list by taxa abundance and specificity (see **Methods**), we compiled a set of marker taxa, designating the water masses in the St. Anna Trough region (**Figure 5D**). The surface KC water mass was specifically enriched with AEGEAN-169 marine group (reaching 2.17%) and SAR11 clade IV (up to 2.25%), while the MW mass was marked with *Planktomarina* (9.95%) and *Sulfitobacter* (6.69%). The BSBW water mass was enriched with *Nitrincolaceae* and *Amylibacter*, where these taxa reached an abundance of 33.1% and 8.02%, respectively. The subsurface CDW was designated by SUP05 cluster (up to 12.2%) and SAR324 clade (6.71%), while CDW+BSBW mass was marked with Chitinophagales (up to 4.72%) and *Flavobacteriaceae* (OTU8 was the most specific and reached 11.2%).

We conclude that despite quantitative rather than qualitative differences between water mass microbiomes in the St. Anna Trough region, we revealed their microbial marker taxa.

## Discussion

In this study, we tested whether seawater bacterial communities are indicative of water masses in a hydrologically complex and sparsely sampled Arctic region of the northeastern Barents Sea and the St. Anna Trough. Using hydrographic analysis, we resolved Barents Sea Branch Water (BSBW), Keldysh Current water (KC), Arctic Surface Water (ASW), Meltwater (MW), Cold Dense Water (CDW), and mixed CDW+BSBW in surface and sub-surface layers of the study area. Microbiome clustering largely reproduced this structure, resolving MW-like, KC-like, CDW-like, and CDW+BSBW communities, but not ASW. Water-mass origin alone explained 56.3% of microbiome variation, whereas water-mass origin, temperature, salinity, and depth together explained 61.2% of variation. We further observed that most taxa were present across all water masses, whereas differences between water masses were explained primarily by changes in OTU relative abundances. Together, these results support seawater microbiomes as complementary biological markers of water-mass origin and identify Keldysh Current as previously unnoticed contributor to “biological Atlantification” in the Eurasian Arctic.

### Hydrophysical structure and water-mass transformation in the St. Anna Trough region

Our hydrographic analysis of the northeastern Barents Sea and St. Anna Trough revealed Atlantic-derived and locally formed Arctic waters linked by transformation and mixing.

Atlantic-derived waters were represented by BSBW and the recently described Keldysh Current (KC), the latter of which occupied the southeastern and eastern flanks of the trough as a warm, relatively saline surface branch derived from the northern periphery of BSBW (Osadchiev et al., 2026). Farther north, recent sea-ice retreat maintained a MW lens at the surface, while ASW likely persisted below MW and above the underlying dense layer; this intermediate position probably explains why ASW was not fully resolved by the 0 and 30–50 m sampling scheme. In the subsurface layer, BSBW remained evident at the southwestern station S3, while locally formed CDW occupied the northern stations. In the southeastern sector, mixing between CDW and Atlantic-derived waters produced the CDW+BSBW assemblage, which was squeezed between the bottom-advected BSBW layer and the surface-advected KC. The localized deepening of KC near S69, close to its formation region, and the CDW-like character of station S10 close to the presumed CDW formation area further indicate that the study region is best viewed as a transformation corridor rather than as a simple Atlantic-to-Arctic gradient. This interpretation is consistent with the superposition of surface KC, subsurface and bottom BSBW, and locally formed CDW previously described for the Barents-Kara gateway (Osadchiev et al., 2022; Osadchiev et al., 2026; Zabudkina et al., 2025).

### Water-mass origin and transformation shape seawater microbiome structure

Our results demonstrated that microbial community composition closely followed the hydrological structure of the northeastern Barents Sea and the St. Anna Trough. Water-mass origin explained the majority of the variation in bacterial community composition (56.3%), exceeding the explanatory power of salinity, temperature, or depth, considered separately, while the combined model performed marginally better and explained 61.2% of the total variation. Because water-mass identity integrates both physicochemical conditions and formation history, these results indicate that microbiome composition reflects more than a single environmental variable. Similar associations between hydrography and bacterial community structure have been reported in the North Atlantic and across the Arctic Ocean, where Atlantic-derived waters retain characteristic microbial assemblages despite extensive transport and transformation (Galand et al., 2010; Agogué et al., 2011; Han et al., 2015; Vipindas et al., 2023).

Interestingly, all delineated water masses, except ASW, produced distinct microbiome clusters: ASW microbiomes were mainly distributed between MW-like and KC-like communities. This pattern is consistent with the regional hydrography: ASW contributes to KC formation and may be overlain by recently formed MW. The absence of a discreet ASW microbiome cluster therefore may reflect the transient and transitional nature of these waters.

A distinct microbiome of the Keldysh Current is particularly notable. Hydrologically, the Keldysh Current represents a mixture between Atlantic-origin waters and Arctic surface waters that remains confined to the upper layer and is transported rapidly eastward along the continental slope (Osadchiev et al., 2026). This suggests that, despite its mixed origin, the Keldysh Current represents not merely a hydrological feature but also a persistent ecological habitat capable of sustaining a distinct microbial community.

We made several observations indicating that microbiome composition may capture fine details of water mass dynamics, which are not fully resolved by standard thermohaline properties alone. First, the abovementioned discrepancy between ASW and the clustering of associated microbiomes. Second, MW microbiomes from two sampling sites (S21 and S63) clustered with KC-like microbiomes. Third, CDW microbiomes from the other two sampling sites (S13 and S60) clustered with CDW+BSBW-like microbiomes. Such discrepancies may indicate the persistence of microbiome signatures after recent advection events and mesoscale mixing. Because microbial turnover occurs on timescales different from physical mixing, microbiomes may retain information about previous environmental conditions and therefore serve as complementary tracers of water mass history.

More generally, our findings support the concept that Arctic microbial communities are structured primarily by environmental and hydrographic conditions rather than by geographic location alone. The spatial extent of our study area spanned approximately 300 by 300 km, and therefore the observed microbiome differentiation cannot be explained by macroscale geographic separation (Winter et al., 2013). Instead, the distribution of microbial communities closely followed water mass boundaries, consistent with previous observations from both Arctic and global oceans showing that environmental filtering dominates over spatial distance in shaping pelagic microbial biogeography (Galand et al., 2010; Winter et al., 2013).

### Marker taxa reflect water-mass origin

Although the majority of microbial OTUs were shared among water mass microbiomes, differential abundance analysis revealed distinct taxonomic signatures associated with them. This observation indicates that differences between water masses are driven primarily by changes in relative abundance of widespread Atlantic and Arctic taxa rather than by replacement of microbial communities.

#### Keldysh Current (KC) microbiomes

KC microbiomes were enriched in several marker oligotrophic bacterial taxa previously associated with warmer Atlantic water, consistently with the origin of this water mass - SAR11 Clade Ia, *Roseibacillus*, SAR86 clade (OTU14), and AEGEAN-169. SAR11 Clade Ia and *Roseibacillus* increase correlated with late-summer Atlantic inflow in Svalbard fjord time series study (Delpech et al., 2021), SAR86 clade increased during the autumn bloom in north-west Atlantic Ocean (El-Swais et al., 2014) and persisted in oligotrophic warm coastal waters in Mediterranean (Korlević et al., 2022), while AEGEAN-169 marine group also increased in surface Mediterranean water in summer-autumn season (Korlević et al., 2022) and is a signature taxon of Atlantic waters in the Fram Strait (Priest et al., 2023). Interestingly, the AEGEAN-169 marine group was associated with expanding and warming oligotrophic conditions globally (Getz et al., 2023), potentially illustrating the “biological Atlantification” of the Arctic via the KC inflow.

Information on other KC marker taxa – SAR11 clade IV and SV1-3 – is scarce, and they may be explored as novel potential signatures of Atlantic-derived waters in the northeast Barents Sea.

#### Meltwater (MW) microbiomes

MW microbiomes were enriched in marker copiotrophic taxa - *Sulfitobacter*, *Formosa*, *Planktomarina*, OM60 (NOR5), and NS5 marine group (OTU4) - that were previously associated with colder and fresher early-melt-season water, in line with the origin of this water mass. *Sulfitobacter* and *Formosa* abundances positively correlated with early-summer post-bloom water enriched in dissolved organic carbon (DOC) in a Svalbard fjord time series study (Delpech et al., 2021), while *Planktomarina* abundance was positively correlated with DOC and particulate organic carbon (POC) and temperature and negatively correlated with salinity in the North Sea (Giebel et al., 2011). Moreover, many of the MW-associated taxa are known to be associated with phytoplankton blooms. As such, *Sulfitobacter* is associated with *Emiliania huxleyi* blooms (Barak-Gavish et al., 2018), was shown to stimulate the growth of diatom algae (Amin et al., 2015), and has been found increased in eutrophic bacterioplankton communities of the southern Chukchi Sea (Lee et al., 2019). *Formosa* is also enriched in algal blooms (Teeling et al., 2012; Teeling et al., 2016), and has a genome encoding a broad repertoire of polysaccharide utilization loci enabling the degradation of polymer-rich organic matter (Mann et al., 2013). Similarly, OM60 (NOR5) was found enriched in algal blooms and implicated in the utilization of different phytoplankton-derived carbon sources (Teeling et al., 2016; Thiele et al., 2023; Yang et al., 2015). *Planktomarina* is a potential particle-specialized taxa (Delpech et al., 2021; Jain and Krishnan, 2017) involved in decaying phytoplankton blooms (Giebel et al., 2011; Wietz et al., 2024). NS5 marine group is associated with algae blooms and organic matter degradation (Signori et al., 2018), an indicator of eutrophication (Kopprio et al., 2021), with genomes harboring multiple putative glycoside hydrolase genes (Ngugi and Stingl, 2018).

Though the described pattern overall aligns with Svalbard fjord time series data (Delpech et al., 2021), several taxa enriched in MW drop out. As such, *Planktomarina*, OM60 (NOR5) clade, and NS5 marine group were elevated in late-summer water in the Svalbard fjord affected by Atlantic inflow, indicating that a combination of global (e.g., water mass origin) and local (e.g., algal bloom, river runoff) factors shape Arctic microbial communities.

#### Barents Sea Branch Water (BSBW) microbiomes

BSBW microbiomes harbored marker bacterial taxa previously associated with warmer Atlantic-derived waters, matching the origin of this water mass. As such, *Amylibacter* was a signature taxon of the Atlantic water-associated microbiome cluster in the Fram Strait (Priest et al., 2023); its abundance showed significant positive correlation with temperature and salinity in the Barents Sea (Aalto et al., 2022) and increased in the Arctic Ocean over summer (Priest et al., 2025). ML602J-51 taxon abundance increased in coastal Mediterranean waters during the unusually warm year 2016 (Trombetta et al., 2022), and this taxon was also associated with estuarine communities (Kujat et al., 2026). *Nitrincolaceae* (synonym of *Oceanospirillaceae*) abundance increased in June in the Svalbard fjord time series study (Delpech et al., 2021); it was associated with the “bloom” cluster in Fram Strait (Weitz et al., 2024) and found enriched in the northeastern Barents Sea (microbiome cluster-2) (Namsaraev et al., 2023).

#### Cold Dense Water (CDW) microbiomes

The CDW microbiome marker taxa were represented by typical Arctic subsurface groups (SUP05, SAR324, and SAR92 clades) (Thiele et al., 2023; Weitz et al., 2024; Cardozo-Mino et al., 2021), several of which were also associated with algal blooms in the Arctic and northwest Atlantic (SAR92 and SAR86 (OTU23) clades) (El-Swais et al., 2014; Teeling et al., 2016; Thiele et al., 2023; Priest et al., 2025). SUP05 and SAR92 taxa were previously associated with cluster-2 microbiomes of the northeastern Barents Sea (Namsaraev et al., 2023), and, together with SAR324, represent residential signatures of Arctic water masses detected in the Fram Strait (Wietz et al., 2024; Priest et al., 2023).

The overall ecological pattern of these taxa matches well with the Arctic origin of CDW. In conflict with this, SUP05 was also associated with Atlantic waters in the Fram Strait in a different study (Priest et al., 2023), and SAR92 abundance was elevated in late-summer samples from a Svalbard fjord time series study, likely affected by Atlantic inflow (Delpech et al., 2021), while SAR86 clade was found persisting in oligotrophic warm coastal waters in the Mediterranean (Korlević et al., 2022). Such discrepancies may reflect that different members of these taxonomic groups might have different ecology and/or CDW could experience mixing with adjacent water masses – presumably bloom-affected surface MW/ASW and southern, Atlantic-derived, KC/BSBW. The first explanation is supported by the fact that different members of the SAR86 clade were enriched in different water masses in this study – OTU14 marked KC, while OTU23 marked CDW.

#### Mixed CDW+BSBW microbiomes

The CDW+BSBW microbiome was marked with taxa implicated in complex polymer degradation and frequently found in particle-associated microbiomes: *Flavobacteriaceae* (NS9 marine group, NS5 marine group – OTU20, and unassigned OTU8) and Chitinophagales (Martínez-García et al., 2022; Urvoy et al., 2022; Ebihara et al., 2024; Preston et al., 2020). Oceanic fronts were shown to enhance particle aggregation (Stemmann et al., 2008). Considering the mixed origin of the CDW+BSBW, we hypothesize that increased particle formation may be expected in this water mass, which could potentially stimulate the development of particle-associated communities.

Interestingly, these taxa were not positively correlated with *Pseudoalteromonas* abundance (**Supplementary Figures 3**-**4**), suggesting the existence of two distinct particle-associated microbiomes in the study area: CDW+BSBW-associated (Flavobacteriaceae/Chitinophagales) and sporadic and widespread *Pseudoalteromonas*-dominated (*Pseudoalteromonas*/*Paraglaciecola*/*Colwellia*/*Oleispira*/*Psychrobacter*). While Flavobacteriaceae are known degraders of algae-derived polysaccharides (Cai et al., 2026), *Pseudoalteromonas*, *Colwellia*, and *Glaciecola*/*Paraglaciecola* were identified in copepod-associated communities (Holmström, Kjelleberg, 1999; Fontanez et al., 2015; Smriga et al., 2016; Moisander et al., 2015; Heins et al., 2021). Based on that, we hypothesize that these microbiomes may represent, respectively, phytoplankton-associated and zooplankton-associated communities, but further studies focused on collecting and profiling of different particle types are needed for the validation of this hypothesis.

Taken together, we conclude that the known ecology of the detected marker taxa is consistent with the identified water-mass origin. Specifically, our observations are in line with a model that KC harbors an oligotrophic community, influenced by Atlantic-derived taxa associated with warmer and more saline water, while MW represents a copiotrophic post-bloom community more typical for colder and fresher seawater. The bloom effect can also explain why ASW lacks a distinct microbiome, which may be hidden by bloom-associated shifts in the bacterial community; however, the lack of direct phytoplankton measurements in this study leaves this as a hypothesis. BSBW community composition is associated with warmer Atlantic-derived waters, while CDW is enriched in resident Arctic subsurface groups, and mixed CDW+BSBW is influenced by particle-associated microbiomes.

### Implications for “Atlantification” and “biological Atlantification” of the Eurasian Arctic

Enrichment of the signature Atlantic-associated taxa within distinct KC and BSBW microbiomes provide new evidence that “Atlantification” of the Eurasian Arctic is not limited to the transport of heat, salt, and nutrients but also involves “biological Atlantification” of microbial communities. Our results indicate that not only deep Atlantic layers but also transformed Atlantic waters in the upper 50 m transported eastward along the continental slope and through the St. Anna Trough contribute to this process. Therefore, we highlight the Keldysh Current as an important biological corridor potentially facilitating the eastward expansion of Atlantic-associated microorganisms into the Eastern and Central Arctic.

Our observations complement the concept of “biological Atlantification” previously described for macrofauna, phytoplankton, and zooplankton communities in the Arctic Ocean (Oziel et al., 2020; Ingvaldsen et al., 2021; Wold et al., 2023). Recent studies in the Fram Strait and Eurasian Basin have demonstrated that increasing Atlantic water influence leads to shifts in microbial community composition toward Atlantic-type assemblages (Priest at al., 2023; Wietz et al., 2024). Our results indicate that similar processes are occurring in the northeastern Barents Sea and the St. Anna Trough, where Atlantic-derived microbial signatures remain detectable even after substantial physical transformation of the original water mass. The persistence of KC-associated taxa such as SAR11 clades Ia and IV and AEGEAN-169 suggests that biological signals may survive longer than conservative hydrographic tracers and therefore provide an additional indicator of Atlantic influence. As “Atlantification” continues to intensify in the Eurasian Arctic, the transport of microbial communities by currents such as the Keldysh Current may increasingly contribute to modifications of regional biogeochemical cycling, organic matter processing, and deeply affect native ecosystems. Long-term monitoring of microbial communities could therefore provide a sensitive tool for detecting ongoing ecological consequences of Arctic climate change and the progressive expansion of Atlantic influence into the Arctic Ocean, and our study established the foundation for such monitoring.

## Acknowledgements and Funding

The sampling was performed during an expedition within the framework of the Russian national scientific and educational program Floating University (research project no. 075-03-2024-117). This research was funded by the Moscow Institute of Physics and Technology under the Agreement No. 075-03-2026-305 dated January 16, 2026 (processing of hydrographic data); the state assignment of Shirshov Institute of Oceanology No. FMWE-2024-0015 (analysis of hydrographic data).

## Data availability statement

The raw sequencing data are publicly available in the NCBI Sequence Read Archive (SRA) under the Project Accession number PRJNA1359548. The scripts used for data analyses and visualization, and the source data for SpeSpeNet correlation networks, are available on GitHub (https://github.com/chorzow/16S_PE/ and https://github.com/rusanovaA/Marine-water-analyses).

## Author contributions

OK: Methodology, Investigation, Writing – original draft, Writing – review and editing, Conceptualization, Funding acquisition; IPD: Visualization, Methodology, Investigation, Writing – original draft; AO: Methodology, Investigation, Writing – original draft, Writing – review and editing; AR: Visualization, Methodology, Investigation, Software; EC: Methodology; DaSi: Methodology; DaSl: Methodology; AS: Visualization, Investigation, Software, Writing – review and editing; ZZ: Methodology; AD: Methodology; AI: Methodology, Resources, Funding acquisition; DmSu: Writing – original draft, Writing – review and editing, Investigation, Visualization, Methodology, Conceptualization, Funding acquisition.

