## Supplementary Figures for "Water-mass origin and transformation shapes seawater microbiomes in the northeastern Barents Sea and the St. Anna Trough"

**Supplementary Materials**

**Supplementary Figures**

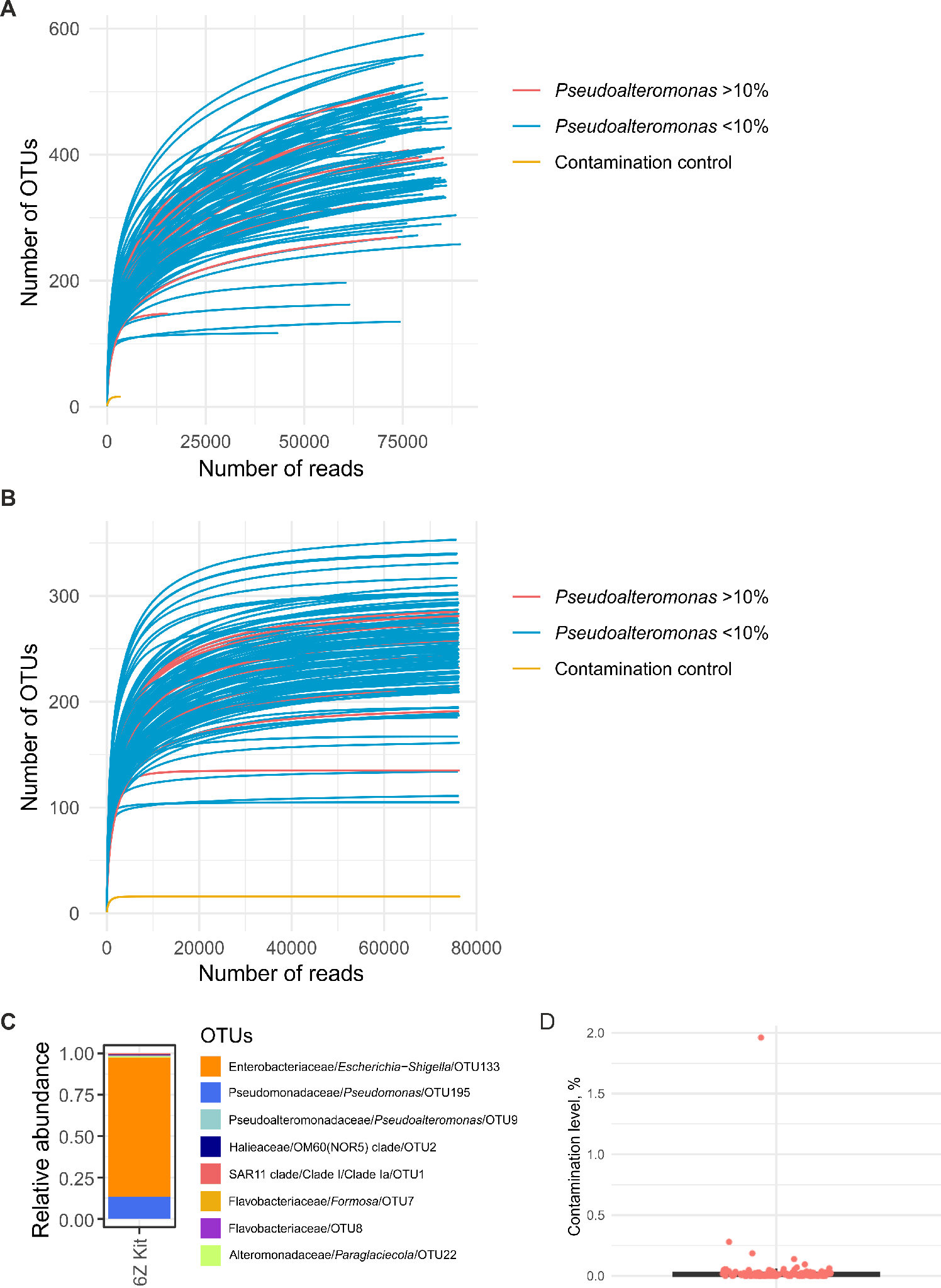

**Supplementary Figure 1**. Rarefaction curves obtained for 16S rRNA gene amplicon sequencing datasets. (**A**) Rarefaction curves for initial datasets before data normalization and removal of sparse OTUs. (**B**) Rarefaction curves after data normalization and removal of sparse OTUs. Samples with a high relative abundance (≥10%) of *Pseudoalteromonas* OTUs marked in red; contamination control shown in yellow; other samples shown in blue. (**C**) Relative abundance barplot for the contamination control sample, profiled with 16S rRNA gene metabarcoding, at the OTU level. (**D**) Boxplot representing contamination levels of seawater samples profiled with 16S rRNA gene metabarcoding. The contamination level was defined as a cumulative fraction of OTUs per sample, which are also present in the laboratory control at a relative abundance of >10%.

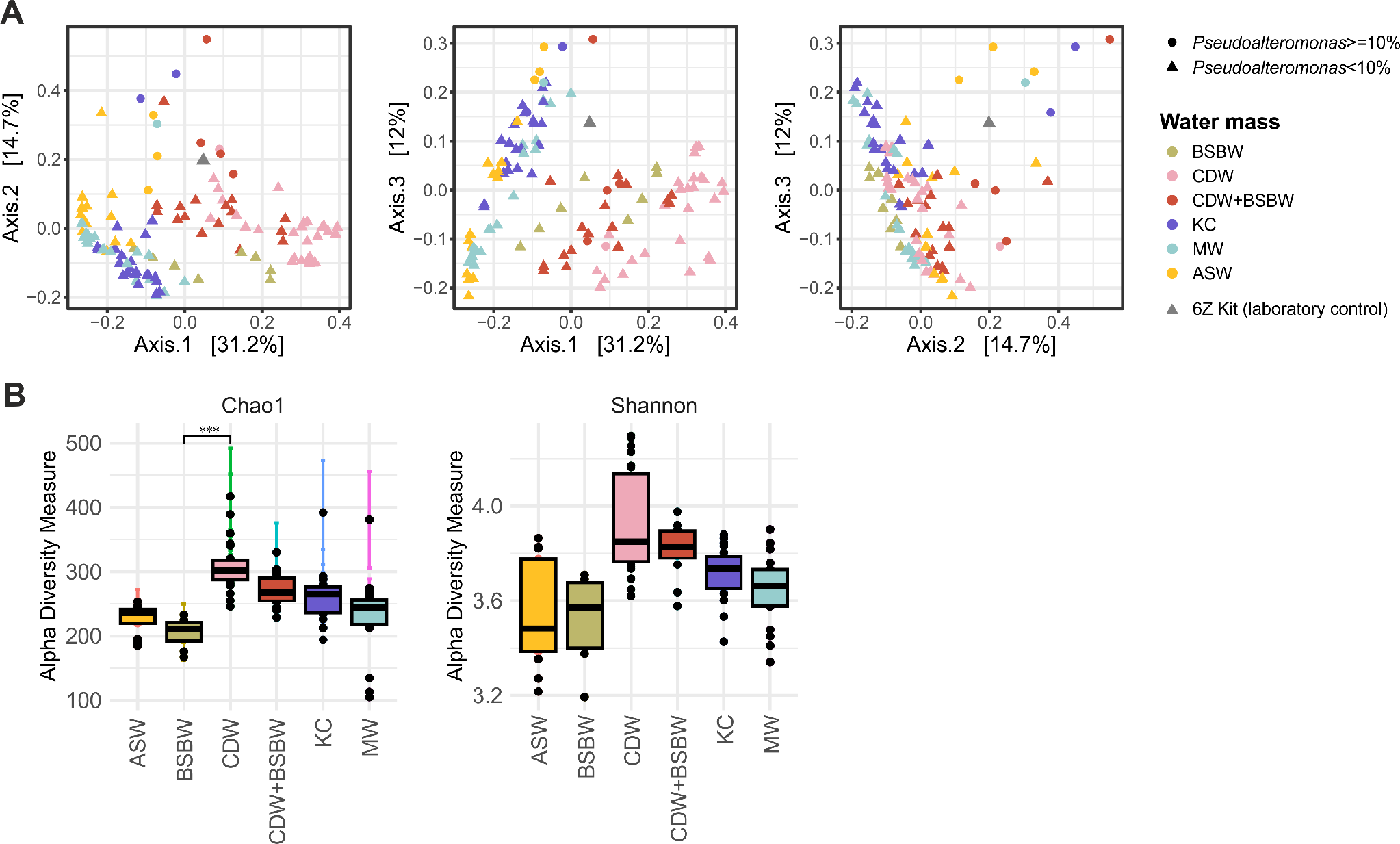

**Supplementary Figure 2**. Beta- (PCoA based on Bray-Curtis dissimilarity) and alpha- diversity (Chao1 and Shannon indices) for collected seawater samples. (**A**) Data for all microbiome samples. Three PCoA projections are shown from left to right. Samples with high (≥10%) and low (<10%) relative abundance of *Pseudoalteromonas* OTUs are marked with dots and triangles, respectively. Samples are colored according to the water mass of origin. (**B**) Alpha-diversity for microbiome samples obtained from different water masses. Data for microbiome samples with low (<10%) relative abundance of *Pseudoalteromonas* OTUs in considered. Samples are colored according to the water mass of origin.

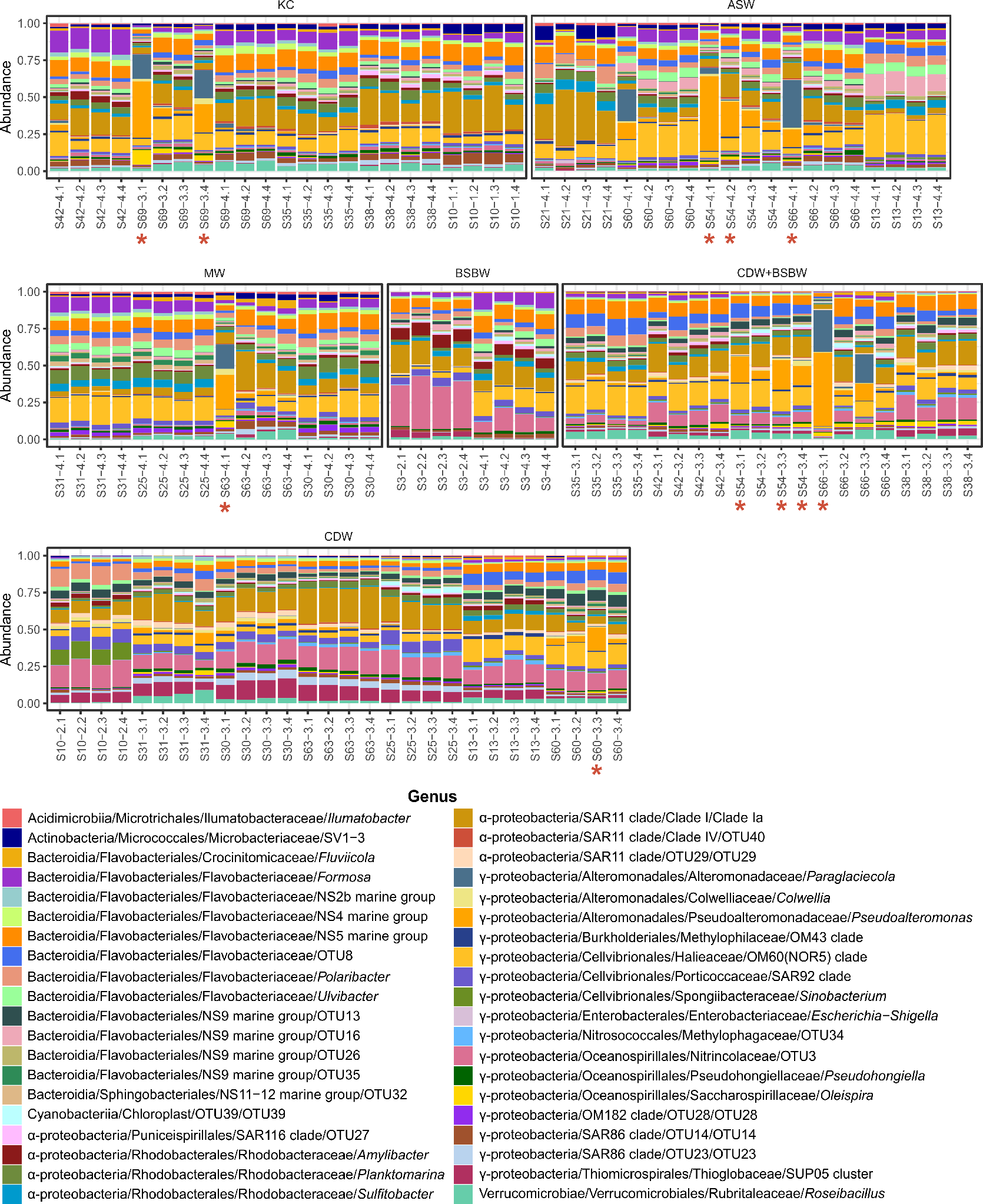

**Supplementary Figure 3**. Relative abundance barplots for all 115 seawater samples, collected and profiled with 16S rRNA gene metabarcoding. Samples are grouped by water masses. Samples with a high relative abundance of *Pseudoalteromonas* OTUs (≥10%) are marked with red asterisks. Data are shown for the top-40 genera sorted by average relative abundance. If a genus was not determined, an OTU ID is reported.

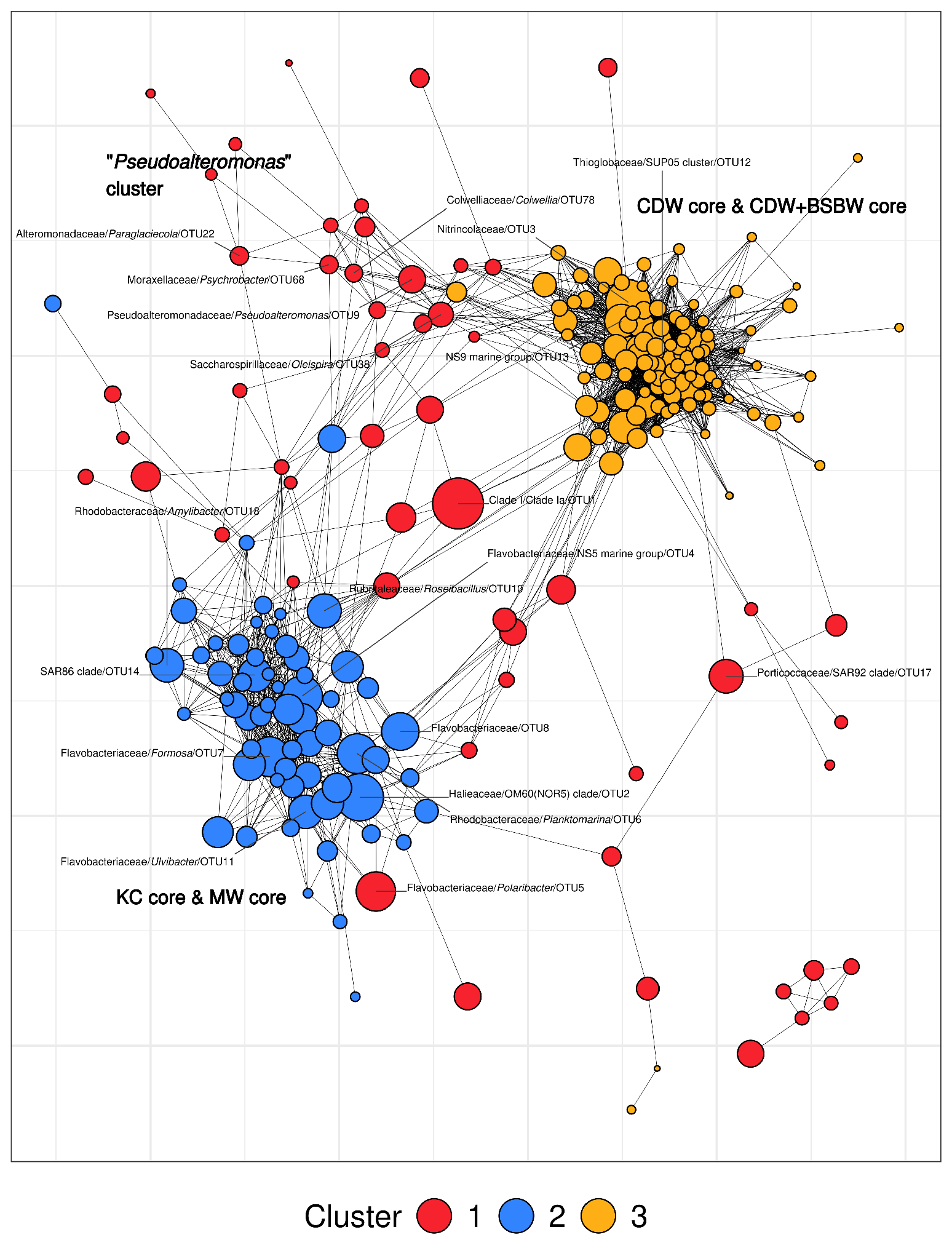

**Supplementary Figure 4**. SpeSpeNet correlation network constructed from core water mass samples. The network includes OTUs with an occurrence threshold of 5 and a relative abundance threshold of 0.1%. Spearman correlations with a coefficient ≥ 0.5 were used to define network edges. Clusters were identified using the k-means method (*k*=3). Differentially abundant and marker bacterial OTUs are indicated.

**Supplementary Tables**

**Supplementary Table 1**: Description of samples for 16S rRNA gene amplicon sequencing (sample metadata).

**Supplementary Table 2**: OTU counts, sequences, and taxonomy.

**Supplementary Table 3**: SpeSpeNet node description for a network constructed on all core water mass samples.

**Supplementary Table 4**: SpeSpeNet node description for a network constructed on surface core water mass samples.

**Supplementary Table 5**: SpeSpeNet node description for a network constructed on subsurface core water mass samples.
